# Improving generalizability for MHC-I binding peptide predictions through geometric deep learning

**DOI:** 10.1101/2023.12.04.569776

**Authors:** Dario F. Marzella, Giulia Crocioni, Tadija Radusinović, Daniil Lepikhov, Heleen Severin, Dani L. Bodor, Daniel T. Rademaker, ChiaYu Lin, Sonja Georgievska, Nicolas Renaud, Amy L. Kessler, Pablo Lopez-Tarifa, Sonja I. Buschow, Erik Bekkers, Li C. Xue

## Abstract

The interaction between peptides and major histocompatibility complex (MHC) molecules is pivotal in autoimmunity, pathogen recognition and tumor immunity. Recent advances in cancer immunotherapies demand for more accurate computational prediction of MHC-bound peptides. We address the generalizability challenge of MHC-bound peptide predictions, revealing limitations in current sequence-based approaches. Our structure-based methods leveraging geometric deep learning (GDL) demonstrated promising improvement in generalizability across unseen MHC alleles. Further, we tackle data efficiency by introducing a self-supervised learning approach on structures (3D-SSL). Without being exposed to any binding affinity data, our 3D-SSL outperforms sequence-based methods trained on ∼90 times more datapoints. Finally, we demonstrate the resilience of structure-based GDL methods to biases in binding data on an Hepatitis B virus vaccine immunopeptidomics case study. This proof-of-concept study highlights structure-based methods’ potential to enhance generalizability and data efficiency, with important implications for data-intensive fields like T-cell receptor specificity predictions, paving the way for enhanced comprehension and manipulation of immune responses.

## Introduction

Peptide-major histocompatibility complex (MHC) interactions play a key role in the immune surveillance system, as T cell discrimination between self and non-self relies on T cell receptor (TCR) recognition of peptides presented by Major Histocompatibility Complex (MHC) molecules. The accurate identification of peptides presented by MHC on the cell surface is crucial for understanding autoimmune diseases^1^, recognizing pathogens^2^, and addressing transplant rejection^3^. The recent notable clinical advancements in cancer immunotherapies^4,5^, specifically targeting tumor-associated or tumor-specific peptide-MHC complexes, underscore the urgency to advance computational methods for identifying MHC-bound peptides^6,7^. With over 14,000 human MHC-I proteins^8^ encoded by the canonical HLA genes (HLA-A, HLA-B, and HLA-C), the theoretical 20^9^ 9-residue peptides (called 9-mers) create impractical *in vitro* testing scenarios. This calls for the development of *in silico* peptide-MHC (pMHC) binding prediction methods.

Much effort has been devoted to design predictors for pMHC binding^9^. Most state-of-the-art (SOTA) predictors are sequence-based machine learning methods^10–12^ trained on a large amount of experimental binding data for pMHC (∼600K for MHC-I^11^ and >480K for MHC-II^10^). Sequence-based (SeqB) methods take amino acid sequences as input to predict binding affinity (BA) or eluted ligand rank. In addition to this, NetMHCpan uses a structure-derived concept like the MHC pseudosequence^13^, utilizing only the MHC residues observed to interact with the peptide in X-ray structures instead of the whole MHC sequence. MHCflurry 2.0 adds to the MHC-I binding prediction an antigen processing model, integrating both scores in its final MHC presentation predictor. These methods are reported to perform well on most of 214 well-studied alleles with hundreds of thousands of mass spectrometry data.

While these approaches have shown considerable advancement and contributions to clinical trials^14,15^, they are not without limitations and have been shown to provide highly discordant predictions^16,17^. A notable constraint stems from their reliance on extensive data, posing challenges for the thousands of less-explored HLA alleles with limited available binding information. Moreover, redundancy reduction strategies, such as the removal of identical peptides or using independent test sets^11,12,18^, are not consistently implemented. This lack of emphasis on ensuring dissimilarity between test and training data can contribute to an overly optimistic performance. Finally, these methods typically train and test on a small subset of MHC alleles, which is a limiting factor considering the broad spectrum of human alleles. Consequently, predictions for underrepresented alleles in these sets may exhibit lower accuracy, akin to an out-of-distribution (OOD) issue in machine learning, where real-case scenarios differ significantly from training data, introducing challenges in generalization^19–21^.

A structure-based (StrB) method can have several compelling advantages over SeqB methods^22^ as it can 1) integrate the huge amount of experimental pMHC binding affinity data with physico-chemical properties of 3D interfaces, 2) reflect minor changes of mutations in 3D space and energy landscape, and 3) naturally handle variable peptide lengths in 3D space. The booming advances of geometric deep learning (GDL)^23–26^, a specialized branch tailored for 3D objects, underscore the timely opportunity for enhancing pMHC binding predictions through the development of StrB methods. Importantly, the highly conserved MHC structures observed across various species, together with the constrained peptide binding within the MHC binding groove, establish a foundation for constructing accurate pMHC 3D models^27,28^. These models, in turn, serve as reliable input data for GDL algorithms.

Much efforts have been devoted to design StrB methods for predicting MHC-binding peptides with different degrees of success in the past decades^22^. Typically, energy terms, statistical potentials and structural descriptors (e.g., the number of polar-polar interactions) are used as input of machine learning algorithms^22^. Based on 77,000 3D models generated with APE-Gen^28^, the authors used Rosetta^29^ to extrapolate energy scores at each peptide position and provided these scores as vectors to train a random forest predictor. Recently Wilson *et al*.^30^ have developed a StrB binding motif prediction method, for which they trained inception convolutional neural networks (CNN) on electrostatic potentials of MHC structures, demonstrating high prediction speed and precision. And MHCfold predicts pMHC 3D structures and the interactions at the same time and achieved promising results in both tasks^31^.

In this study, we develop end-to-end GDL methods that directly analyze structural features at atom or residue level to predict the binding between peptide and MHC. We set out to demonstrate the generalizability issue for pMHC binding predictions and evaluate whether our GDL-based StrB methods have a more robust generalization than SeqB methods for pMHC binding predictions. We compared three different GDL StrB networks with two SeqB networks on an allele-clustered scenario, deliberately minimizing the similarity between the test and training data. To assess the models’ performance, we juxtaposed these results with those obtained from a shuffled dataset, enabling an evaluation of the accuracy drop when extrapolating to unseen cases. In addition, being inspired by the success of the self-supervised learning (SSL) in natural language processing^32,33^, we introduced a novel self-supervised GDL method, 3D-SSL. Remarkably, this method showed superior potential in data efficiency even without exposure to any binding affinity data. Finally, we demonstrate the robustness of StrB methods against biases in the binding data through a study-case on HBV vaccine design. This study underscores the potential strength derived from the integration of 3D physics-based modeling and data-driven geometric deep learning to enhance both generalizability and data efficiency. These findings hold broad implications for the fields of immunology and therapy design.

## Results

### Data separations using allele clustering

We set out to test the generalization power of StrB over SeqB approaches to unseen data through two different data splitting configurations of the same binding affinity dataset^11^: a randomly shuffled one, resembling SOTA-used datasets, and an allele-clustered one. The allele-clustered data configuration was designed to evaluate the applicability of the neural networks to a test set composed of unseen data, the distribution of which might deviate significantly from the training data (e.g. patients with less-studied alleles).

In the shuffled configuration, we randomly split the data in training, validation and test sets, using target stratification to ensure similar target distribution in all data sets (**Fig. 1A**). This configuration aligns with common practices found in existing literature, where the test set is typically either randomly extracted from the initial dataset or is an independent dataset sharing similar alleles and peptides distributions^11,18^. In the second experiment (allele-clustered), we clustered the data based on the MHC pseudosequence^13^ using a PAM30-based^34^ hierarchical clustering for each gene, i.e., HLA-A, -B and -C, respectively (**Fig. 1B-D**, see also Methods). Then, from a total of ∼100K binding affinity data (see details in Methods), for each gene we selected the most distant clusters based on the MHC pseudosequence’s dendrogram so that about 10% of each gene’s data (10% for HLA-A, 11% for -B and 12% for -C, respectively) was selected as test set. We split the remaining data into training and validation sets in a target-stratified manner. This design allows the networks to sample a large number of cases from each HLA gene, while strategically incorporating only distant clusters of alleles into the test data. This configuration aims at simulating real-case scenarios in which the network has to provide a prediction on alleles it has never seen or seen infrequently. We build 3D models for all these binding affinity data using PANDORA^35^, which served as input of our GDL approaches (**Fig. 1E**).

**Figure 1.**
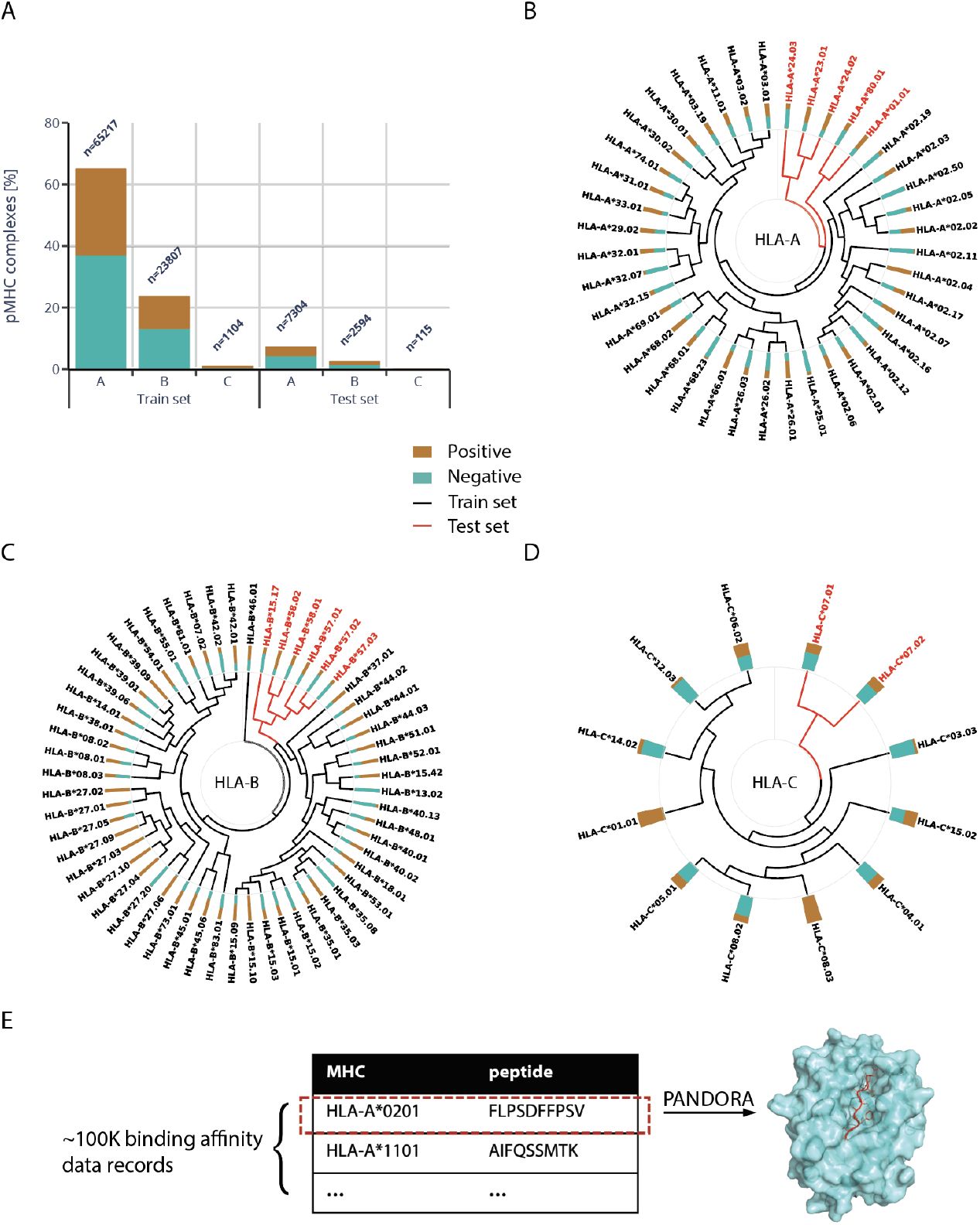
Data overview. A) Shuffled data distribution representation between HLA-A, -B and -C for the training and test sets. B-D) show the hierarchical clustering of HLA-A, -B, -C pseudosequences, respectively, with training data in black and test data in red. In all panels, the mustard/teal bar by each gene label represents the binders/non-binders (positive/negative) ratio for the gene. E) Data enrichment through 3D modeling. We used PANDORA^35^ to generate 20 3D models of each of the 100 178 pMHC data points, thus enriching the sequence information with physics-derived information, such as geometry and physico-chemical features.

### StrB predictors demonstrate greater generalizability than SeqB on distant alleles

For StrB predictors we used different network architectures with different types of physico-chemical features (**Fig. 2A**). We developed an atom-level 3D-CNN and a residue-level GNN using different structural and energy-related features (**Tables 1** and **2**), and a residue-level EGNN using as input features only the residues identities and their xyz coordinates (**Table 3**). 3D-CNNs analyze structural features that are mapped on a 3D grid, capturing local geometric relationships effectively. GNNs and EGNNs treat structures as connected nodes (representing atoms and/or residues) on a graph, updating the features of a node iteratively through passing information of its neighboring nodes^36^. In handling 3D structures, a key requirement for GDL is rotation-translation (RT) invariance, meaning consistent predictions regardless of input orientation. 3D-CNNs capture local geometric relationships but may be sensitive to rotations. GNNs ensure constant outputs even with input rotations. EGNNs further enhance its networks by enabling both the output and all of its features of hidden layers to consistently rotate as well. EGNNs, known for simplicity, are widely used in predicting molecule properties and generating realistic small molecules^24,26,37^. Since many SOTA SeqB predictors are not available to be re-trained for a fair comparison, we developed a Multi-Layer Perceptron (MLP) to represent a baseline sequence-based approach. We also re-trained MHCflurry2.0, being a SOTA architecture available for re-training.

**Table 1.**
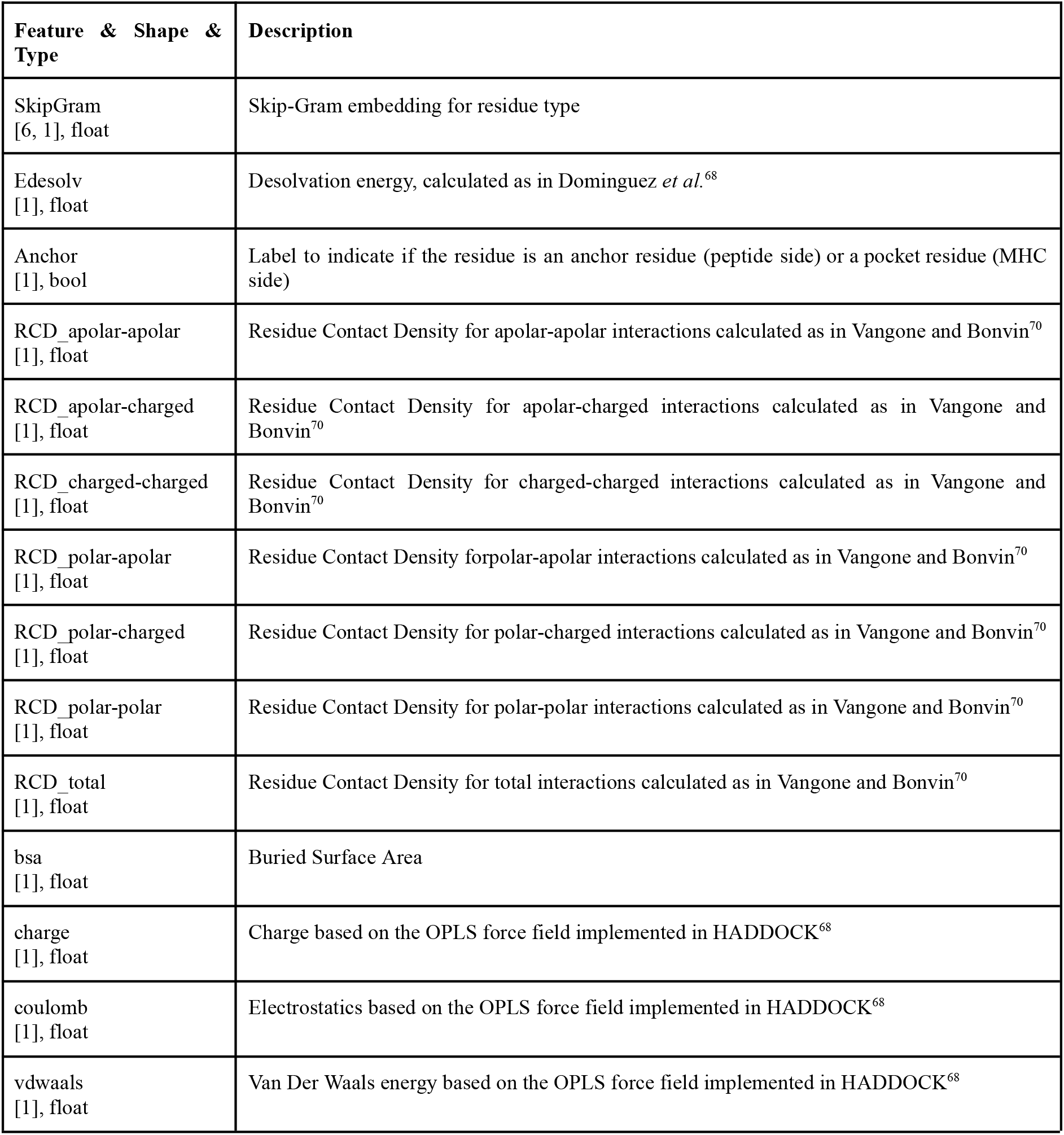
Input features to the CNN architecture. Full features details on built-in deeprank features can be found on https://deeprank.readthedocs.io/en/latest/deeprank.features.html.

**Figure 2.**
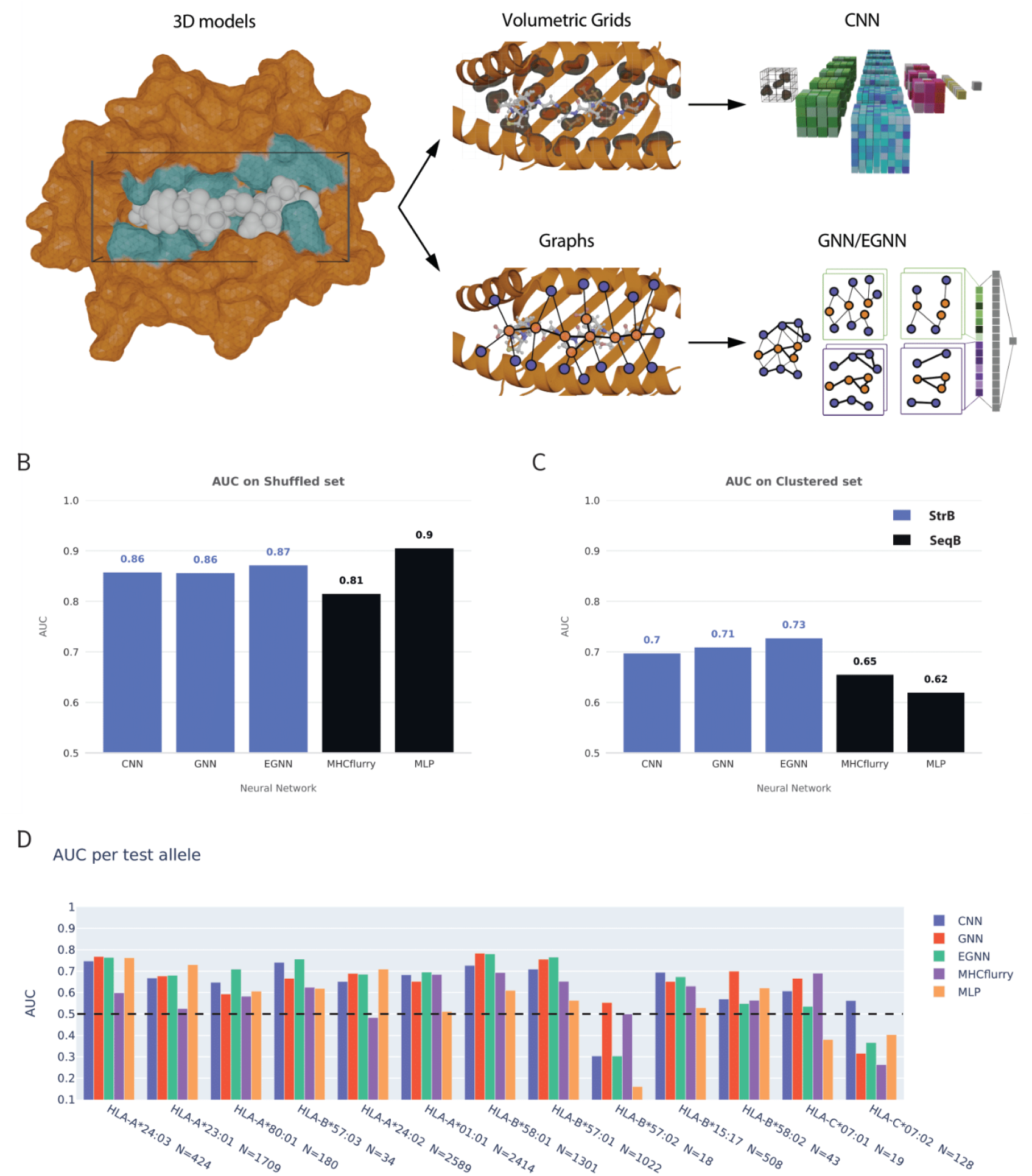
Our supervised StrB methods overview and performances. A) Overview of the pipeline employed for processing pMHC complexes and running the supervised structure-based networks. The process involves: 1) Identifying interface atoms and residues from the pMHC 3D models, 2) converting each pMHC interface into volumetric grids and graphs enriched with geometric and physico-chemical information (**Tables 1-3**), 3) Run the networks on these representations: volumetric grids for the CNN and graphs for GNN and EGNN. GNN and EGNN use similar graph topology, differing on features and message passing framework (see details in Methods). B-C) The performance of StrB and SeqB methods on the shuffled test dataset and on the allele-clustered test dataset, respectively. D) AUC per allele on the allele-clustered test set. Allele name and number of test cases are reported on the x-axis, and the alleles are sorted by sequence distance with the training set. The black dashed line marks the random predictor AUC value of 0.5.

As expected, in the shuffled data configuration, all methods perform well (**Fig. 2B**), with the MLP and our EGNN showing a slight edge over the others, with Area under the ROC Curve (AUC) of 0.91 and 0.87, respectively (see AUCPR details in **Suppl. Fig. 1**). SeqB methods learn the features distribution and can provide good predictions, as suggested from their good performance shown in **Fig. 2B**. StrB methods’ high performance suggests their potential in reaching performance comparable to the current SeqB literature.

In the allele-clustered configuration (**Fig. 2C**), all predictors show a decline in performance, highlighting the generalizability issue. StrB approaches demonstrated more robust performance than SeqB methods on distant alleles. StrB consistently outperforms SeqB by 5-11% in terms of AUC. The substantial performance drop of SeqB methods underscores their susceptibility to the out-of-distribution issue, highlighting the importance of rigorous evaluation for obtaining real-life estimations of predictive methods. MHCFlurry is an ensemble predictor using multiple neural networks, demonstrating better generalization than a basic MLP. **Fig. 2D** shows the performances on each test allele ordered with increasing distance to the training alleles. Computational efficiency for the StrB methods is reported in **Suppl. Table 1**.

Notably, EGNN, our simplest GDL method and the top-performing StrB, achieves a significantly better AUC and AUCPR compared to SeqB methods by 8-11% (**Fig. 2C, Suppl. Fig. 1**). EGNN, using only amino acid types and distances, virtually distinguishes itself from the SeqB methods solely by integrating the spatial relative position of residues and the network’s treatment of this information. This suggests that the enhanced generalization ability of our StrB methods relies on the inclusion of geometrical information in the 3D models and the GDL’s capacity to incorporate such details in the learning process, rather than being dependent on a specific set of features or network architecture.

### Self-supervised learning demonstrates superior data efficiency

Self-supervised learning is a game-changing technique for natural language processing^32,33^. Many well-known architectures, including BERT^32^, GPT-X^33^, MAEs (Masked Autoencoders)^38^ are SSL at their core. Unlike supervised prediction, where a network is trained to directly predict a target value (a binding label, in our case), in SSL we mask or noise part of the data, and task the network with unmasking/denoising the input. Because of this self-prediction nature, SSL is potentially data efficient. Here we extend SSL to 3D and evaluate its performance on pMHC binding predictions.

#### Training

Our 3D-SSL method is trained on a masked residue prediction (MRP) task (**Suppl. Fig. 2**). At training time, 20% of the residues in the pMHC complex is masked, i.e. set to a dedicated ‘unknown’ token. An EGNN is trained to predict the residue types of the masked nodes. In essence, the MRP pushes the network to learn which residues are likely to be found in a given environment, effectively learning a multi-body statistical potential^39–41^. For proof-of-concept, our 3D-SSL is currently trained only on 1,245 X-ray structures from TCRpMHC complexes (**Fig. 3DD**, Methods).

**Figure 3.**
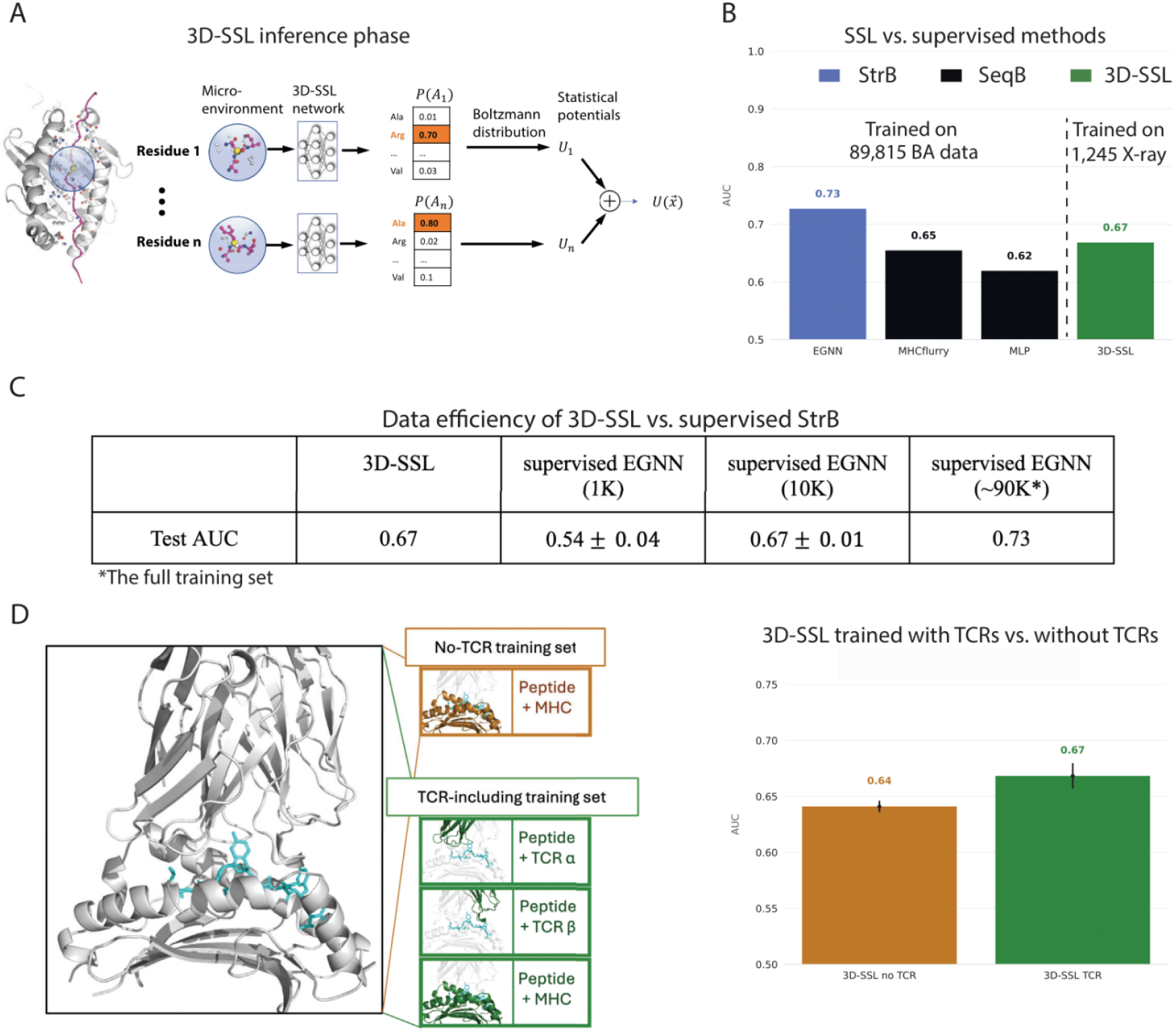
Self-Supervised learning approach (3D-SSL) overview and performances. A) the inference stage of 3D-SSL. 3D-SSL is an EGNN network. It takes the microenvironment of a masked peptide residue as input, predicts the probability of amino acid identities of the masked residue, then converts the probability into statistical potential using Boltzmann distribution as the energy contribution of this microenvironment. For the training stage of 3D-SSL, see **Suppl. Fig. 2**. B) Comparison of 3D-SSL (i.e., unsupervised EGNN) with the same EGNN network trained in a supervised manner and with SeqB approaches, on the allele-clustered dataset. There are no allele overlaps between the 3D-SSL training set and the allele-clustered test set. C) Data efficiency of 3D-SSL against supervised EGNN, our top StrB method. The same EGNN architecture is used by 3D-SSL and supervised EGNN (see Supplementary Note 2). We evaluate the effect of training data size (1K, 10K, ∼90K) on supervised EGNN performance. The allele-clustered BA dataset is randomly sampled for training the supervised EGNN. AUCs are averaged over 5 runs to account for the potentially unrepresentative subsets. D) 3D-SSL performance in terms of AUC when trained with and without peptide-TCR structures, as schematized on the left. We repeated the experiments five times to be sure the difference between training with TCRs and without TCRs is not caused by randomness (error bars for standard deviations shown in black).

#### Inference

Once the network has been trained on X-ray structures, we use it to predict the peptide binding of our test allele-clustered dataset. The trained network takes a pMHC 3D model as input, masks each peptide residue and predicts the probability of the 20 amino acid types at the masked position. Innovatively, we convert the probabilities into a statistical potential using a Boltzmann distribution. The 3D-SSL score is a sum of energy contributions from individual peptide residue to the binding (**Fig. 3A**, Methods). We used this final score as a label for the binding affinity.

Our 3D-SSL is surprisingly an efficient learner. On the allele-clustered dataset, 3D-SSL, trained on training on only ∼1K experimental complexes and not being exposed to the binding affinity values, already outperformed the SeqB supervised methods trained on >90K binding affinity data (**Fig. 3B**). This result shows the superior data efficiency potential of SSL networks.

To further estimate the data efficiency of 3D-SSL against StrB methods, we train supervised EGNN on small subsets of the BA training set to estimate the amount of BA data at which supervision becomes more useful than SSL. When comparing the supervised EGNN against 3D-SSL on the allele-clustered set (**Fig. 3C**) we can see the EGNN requires approximately 10,000 BA data points to consistently outperform 3D-SSL, which achieves comparable performance with training on only 1245 data points, roughly ten times less data. This indicates that if many high-quality pMHC structures with BA measurements are available for a given problem, supervised StrB methods will likely result in better predictive performance. However, in a low-data regime, e.g., working with rare alleles or novel neoantigens, training self-supervised models could likely perform better than supervised ones.

Interestingly, we noticed that including peptide-TCR structures in the training set of 3D-SSL improved the prediction for pMHC binding affinity. We have tested two different training sets for 3D-SSL: one including only pMHC structures and one including also peptide-TCR (T-Cell Receptor) structures, extracted from full TCR-peptide-MHC X-ray complexes. We found out that the larger dataset including peptide-TCR structures provided higher binding affinity prediction performance, from AUC of 0.640±0.011 to 0.668±0.005 (**Fig. 3D**). The improved results with peptide-TCR data might indicate that adding competing binding environments contributes to the prediction of the binding affinity between peptide and MHC.

### A case study: Vaccine for chronic Hepatitis B virus (HBV) infection

We compared one of our StrB methods, GNN, to two SOTA softwares on a real-case scenario: an HBV vaccine design study^42^. In this study, HLA-A*02:01 matched dendritic cells (DCs) from healthy donors were exposed to synthetic long peptides containing HBV sequences in order to test which peptides would be presented, and therefore would serve as possible effective vaccine candidates (**Fig. 4A**). Notably, two closely related peptides (HLYSHPIIL, denoted as HL, and HLYSHPIIL**G**, denoted as HG, differing only in an additional C-terminal glycine in HG) were experimentally confirmed as binders through mass spectrometry and an *in vitro* HLA binding assay^42^. However, only HL but not HG is predicted as a binder by the NetMHCpan4.1b web server and the BA predictor of MHCflurry 2.0’s official release (**Fig. 4B**). The HL peptide has been reported numerous times in literature as an HLA-A02 binder^43–45^, thus it is likely present in the training set of NetMHCpan4.1 and MHCflurry 2.0. In a blind test setting (i.e., we did not know the labels of binding or not beforehand when running our StrB methods), we went on to check whether our StrB methods can accurately identify the HL/HG peptides as binders. We first generated 3D models of the pMHC complex between the HL/HG peptides and HLA-A*0201, and then we used our trained GNN network to make predictions. Our GNN was able to predict both peptides as binders with high confidence scores (**Fig. 4B**).

**Figure 4.**
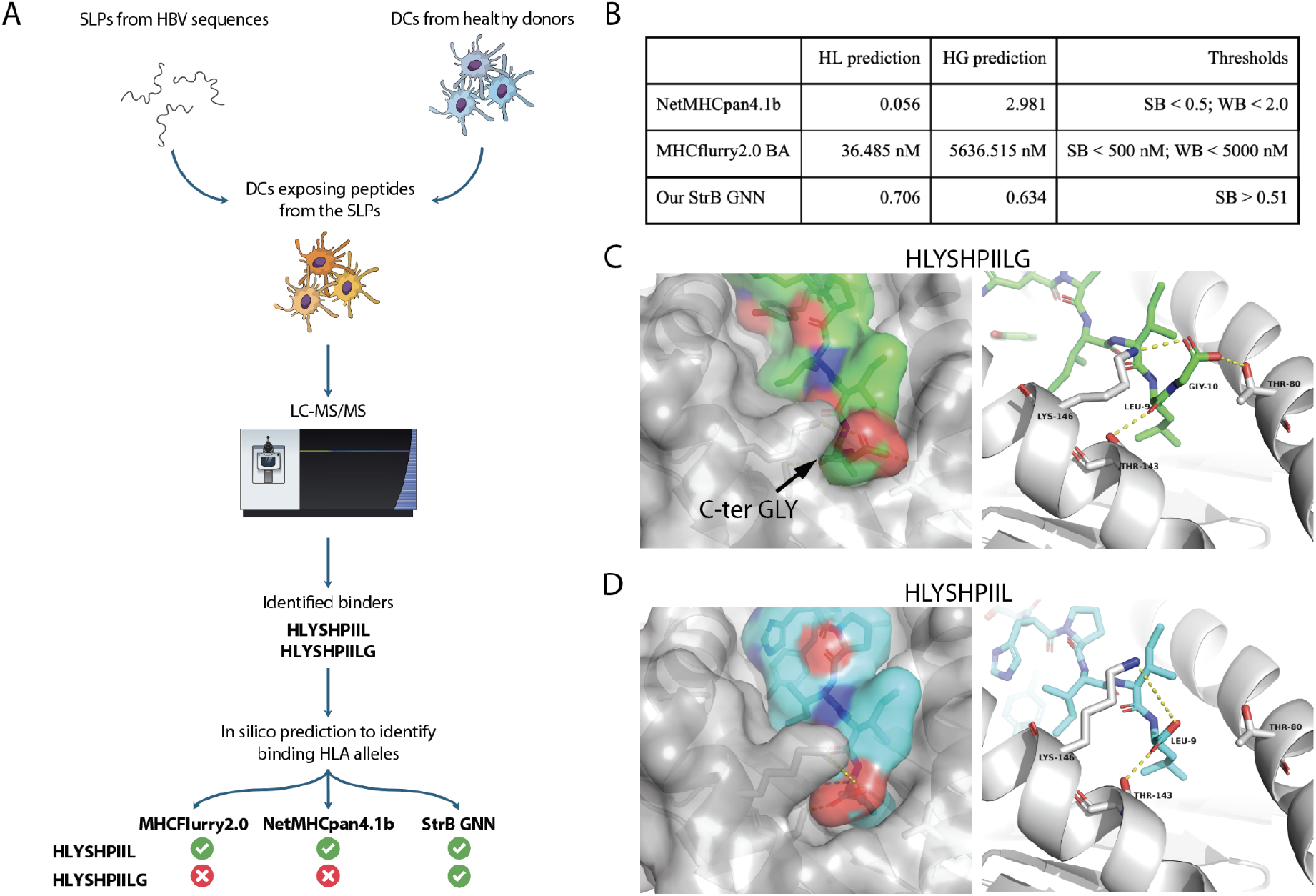
Case study on two peptides in HBV vaccine research. A) HBV case study experiment. Sequences isolated from HBV have been used to synthesize SLPs. The SLPs were then added to DCs cultures from 6 healthy donors. Samples were then analyzed by liquid chromatography tandem mass spectrometry (LC-MS/MS), leading to the identification of peptides presented on the surface of DCs. The peptides were then fed into NetMHCpan4.1b, MHCflurry2.0 BA and our StrB GNN. Only our GNN successfully identified both peptides as binders. B) Outputs and cutoffs from predictors tested. The GNN here is trained on the whole ∼100K binding affinity data. C-D) Structural models for HLYSHPIILG (green) and HLYSHPIIL (cyan). Expected polar interactions and hydrogen bonds are shown as yellow dashed lines. Structural models are generated by PANDORA^35^. Visualizations are done in PYMOL^49^.

When inspecting their 3D models it becomes apparent that the HG/HL case is in fact a relatively easy case for structure-based approaches. The C-terminal glycine is overhanging without causing major unfavorable interactions or clashes (**Fig. 4C**, see also **Suppl. Video 1**). The HL peptide can form a weak salt bridge with MHC LYS146 using its C-terminal carboxyl group. The addition of the glycine in HG causes a bigger entropic effect (i.e. the glycine has to reduce its entropy to fit in the pocket), which can be balanced by the interactions of the glycine’s carboxyl group, which takes a weak salt bridge with LYS146 and an extra hydrogen bond with THR80. Overall, this result indicates that structural modeling adds an important biochemical environment to assist structure-based AI to discover the unconventional HG peptide.

It is critical to emphasize that the biases in SeqB networks might result from training data with potential bias. In fact, the proteasome, which is responsible for peptide generation in non-APC cells, has a low probability of cleaving the C-terminal bond of a glycine^46^. As a result, C-terminal glycines, presumably generated by endosomal proteases in antigen presenting cells (APCs), are poorly represented in training sets which contain mostly peptidomes of parenchymal cells/ non-APCs, likely causing the poor performance of SeqB methods on peptides with C-terminal glycines. MHCflurry 2.0 addresses similar challenges with its antigen preprocessing (AP) module^11^, which could filter out peptides not supposed to be processed by non-APC cells^47,48^, such as peptides containing C-terminal glycines. However, this does not justify the inaccuracies in their BA predictor, which should be able to predict these peptides as binders when presented with them. Overall, the robustness of the StrB GNN might prove effective against other issues caused by the biases that SeqB networks inherit from the data, such as the low accuracy in predicting MHC binding of cysteine-containing peptides, caused by underrepresentation of cysteines in the MS-derived immunopeptidomes used for training.

## Discussion

Generalizability is an important topic in a broad range of studies^50–52^. In this proof-of-concept study, we investigated the generalizability of various StrB approaches in predicting peptide-MHC interactions, a pivotal aspect of immune surveillance and a major bottleneck in the design of cancer vaccines^53–55^ and TCR therapies^56,57^. We showed that our StrB networks have greater generalization power compared to conventionally used SeqB networks when evaluated on unseen MHC alleles. Interestingly, our top StrB method EGNN, utilizing only amino acid types and distances, distinguishes itself from SeqB by incorporating spatial positions. This indicates that the enhanced generalization in our StrB methods relies on GDL’s capacity to integrate 3D geometric information during learning, rather than specific features or network architecture. A similar trend can be seen in our self-supervised approach, 3D-SSL, which uses the same features as our supervised EGNN. Notably, 3D-SSL outperforms sequence-based approaches trained on ∼90 times more data, demonstrating its potential data efficiency, being trained on minimal data and features. The presented HBV test case, although not representing an extensive comparison of SeqB and StrB neural networks biases, exemplifies how StrB methods can better capture the biochemical environment of the pMHC complex, potentially minimizing non-biochemical biases in the data.

We demonstrate the efficiency and feasibility of boosting experimental BA data with physics-derived pMHC 3D models to achieve better predictions of MHC-bound peptides. The inherent conservation of MHC structures makes large-scale physics modeling feasible. These models offer essential complementary information to biochemical binding assays, aiding in training highly generalizable algorithms for the identification of MHC-bound peptides. We anticipate that the fusion of 3D modeling with the expanding capabilities of GDL will soon provide robust and valuable tools for prioritizing targets in cancer immunotherapies, thereby accelerating development timelines and significantly reducing associated costs. Furthermore, MHC structures exhibit strong conservation, with peptides confined to the binding groove, making them an ideal toy model for AI research, particularly in geometric deep learning. The rich experimental binding data, combined with our 3D models, holds potential to advance AI algorithms, providing novel solutions and insights for both the AI and immunology communities.

While our study is on pMHC-I, our StrB methods are directly applicable to pMHC-II complexes, which play a pivotal role in cancer immunotherapy responses^58,59^ but their BA is challenging to predict. StrB methods can naturally handle peptide length variability in 3D space while SeqB methods are inherently limited as they take fixed length input. All together, our StrB approaches, especially, our 3D-SSL approach, may provide a data efficiency solution for data hungry fields, for example, the long-standing challenge of TCR specificity predictions^60^. Expanding the 3D-SSL information source to the Protein Data Bank^61^ structures and AlphaFold2^25,62^ models and integrating supervised methods with our 3D-SSL are potential next steps.

## Methods

### Binding affinity data collection

BA data was collected from O’Donnel *et al*.^11^ through MHCflurry 2.0 ‘‘‘download-fetch’’’ command, which yielded a total of 598,450 pMHC binding affinity measurements. We selected only qualitative essays on human alleles and filtered out data points considered non-exact values (i.e. samples assigned to have a value lower or higher than a certain threshold, with no exact value associated). The final dataset consists of 100,178 data points each containing a peptide sequence, an MHC allele name and their measured BA, using a threshold of 500 nM to separate binders from non-binders, resulting in 44,102 binders and 56,076 non-binders from 110 HLA alleles (see **Fig. 1** and Data & Code Availability).

### 3D modeling with PANDORA

The BA dataset was processed with PANDORA (v2.0beta2) software^35^ to generate 3D models of the pMHC complexes for these binding affinity data. For every case, PANDORA uses NetMHCpan4.1^10^ to predict the peptide’s binding core, using it to select the anchor residues to restraint. The 3D models were generated with PANDORA’s fully flexible mode (i.e., 20 3D models per case, with a restraints flexibility allowed of 0.3Å), with the exception that only the G-domain of the MHC is modeled, while the rest of the MHC α-chain and the β-2 microglobulin are ignored. The top molpdf^63^ scored model was selected as the best model and used as the training, validation and test data.

### Data clusterings

In the shuffled data configuration, we split the data into training (90%) and test (10%) sets after random shuffling, stratifying them on the binary target to make sure the positive and negative cases are equally represented in the different sets. During the training phase, 15% of the training set was used as a validation set.

For the allele clustering, allele pseudosequences were collected from O’Donnel *et al*.^11^. As explained in Hoof *et al*.^13^, these pseudosequences only contain the residues at the MHC positions that have been shown in X-ray structures to interact with the peptide, which makes it so every pseudosequence has the same length and no alignment is needed, as every position correspond to the same physical position on the MHC structure. The pseudosequence were separated per gene and scored against each other using a BLOSUM62 scoring matrix, and the so-obtained evolutionary distances were used to build a hierarchical clustering using the cluster.hierarchy module from scipy^64^ for each HLA gene. From these three clusterings (one per gene), we selected the most distant clusters until reaching the closest possible value to 10% of the data (10% for HLA-A, 11% for HLA-B and 12% for HLA-C) to form the test set. All the other data points were left as a train/validation set.

### Training data for 3D-SSL

We extracted 1245 entries by overlapping PDB IDs containing MHCI from ProtCID with the protein-peptide database Propedia^65^. These Propedia entries are composed of 517 pMHC and 731 TCRα-peptide or TCRβ-peptide structures, separated out from the original PDB structures. We consider two SSL datasets from these entries. First, a reduced dataset of 517 only pMHC structures, and all the extracted entries for the second one. This corresponds to two SSL training scenarios, one where the network sees only pMHC structures, and another where it also sees TCRα/β-peptide structures (**Fig. 3C**).

### 3D-CNN

3D-CNNs are translation invariant but not rotation equivariant. For this reason, we decided to align all the pMHC 3D models to prevent any rotation issue and avoid resorting to data augmentation. In order to generate consistent 3D grids for each 3D model, all the selected 3D models were aligned using GradPose^66^. The principal component analysis (PCA) of the peptides coordinates was then calculated and the X, Y and Z axis of each PDB file were translated to match with the three principal components, to make sure the axis of the grid would sit along the axis of biggest variation in the peptides’ backbone.

For the grids featurization, the PDB files of the aligned pMHC 3D models were then processed with DeepRank software^67^ to generate 3D grids as described in Renaud *et al*.^23^. Each grid was set to have a shape of 35x30x30 Å and a resolution of 1 Å. Only residues of each chain within 8.5Å from the other chain are considered “interface” residues and kept for feature calculation. Features were then generated for each atom position and mapped to the grid points with a Gaussian mapping as described in Renaud *et al*.^23^. In addition to DeepRank’s default features, we added an Energy of Desolvation feature calculated as in Domungiez *et al*.^68^, as the difference in the solvation energy between the bound and unbound complex. We have also replaced DeepRank’s default PSSM feature used for sequence embedding with the lighter skip-gram residue embedding taken from Phloyphisut *et al*.^69^. All the features used are summarized in **Table 1**. The network architecture is described in detail in the **Supplementary Note 2**.

### GNN

The PDB files were converted into protein-protein interfaces (PPI) in the form of residue-level graphs using DeepRank2 package^71,72^. The latter is a Python package inherited from our previous package DeepRank^67^ and that offers a complete framework to learn PPI interface patterns in an end-to-end fashion using GNNs. For a given PDB file representing a pMHC complex, we defined the interface graph as all the residues involved in intermolecular contacts, i.e. the residues of a given chain having a heavy atom within an 15 Å distance cutoff of any heavy atom from the other chain. These contact residues formed the nodes of the graph. Edges were defined between two contact residues from distinct chains presenting a minimal atomic distance smaller than 15 Å. Node and edge residue-level features used are summarized in **Table 2**. The network architecture is described in detail in the **Supplementary Note 2**.

**Table 2.** Input features to the GNN architecture. Full features details can be found on https://deeprank2.readthedocs.io/en/latest/features.html.

| Feature & Shape & Type | Description |
| --- | --- |
| NODE FEATURES |  |
| res_type<br>[21, 1], bool | One-hot representation of the node's residue (20 amino acids + unknown). |
| res_size<br>[1], int | The number of non-hydrogen atoms in the side chain. |
| res_mass<br>[1], float | The average residue mass in Da. |
| res_charge<br>[1], int | The charge of the residue in fully protonated state in Coulomb. |
| res_pI<br>[1], float | The isoelectric point, i.e., the pH at which the molecule has no net electric charge. |
| polarity<br>[4, 1], bool | One-hot representation of the polarity of the amino acid: NONPOLAR, POLAR, NEGATIVE, POSITIVE. |
| hb_donors<br>[1], int | The number of hydrogen bond donor atoms in the residue. |
| hb_acceptors<br>[1], int | The number of hydrogen bond acceptor atoms in the residue. |
| res_depth<br>[1], float | The average distance in Å of the residue to the closest molecule of bulk water, computed via BioPython. |
| hse<br>[3, 1], float | Half sphere exposure, which indicates how buried an amino acid residue is in the biomolecule, computed via BioPython. |
| sasa<br>[1], float | Solvent-accessible surface area, in Å<sup>2</sup>, computed via FreeSASA. |
| bsa<br>[1], float | Buried surface area, which represents the area of the complex interface, in Å<sup>2</sup>, computed via FreeSASA. |
| irc_total<br>[1], int | The number of residues on the other chain that are within a cutoff distance of 5.5 Å. |
| irc_negative_negative,<br>irc_negative_positive,<br>irc_nonpolar_negative,<br>irc_nonpolar_nonpolar,<br>irc_nonpolar_polar,<br>irc_nonpolar_positive,<br>irc_polar_negative,<br>irc_polar_polar,<br>irc_polar_positive,<br>irc_positive_positive<br>[1], int | As above, but for specific residue polarity pairings. |

| EDGE FEATURES |  |
| --- | --- |
| same_chain<br>[1], bool | Boolean indicating whether the edge connects nodes belonging to the same chain (1) or separate chains (0). |
| covalent<br>[1], bool | Boolean indicating whether nodes are covalently bound (1) or not (0). Covalency is not directly assessed, but any edge with a maximum distance of 2.1 Å is considered covalent. |
| electrostatic<br>[1], float | Electrostatic potential (also known as Coulomb potential) between two nodes, calculated using interatomic distances and charges of each atom. |
| vanderwaals<br>[1], float | Van der Waals potential (also known as Lennard-Jones potential) between two nodes, calculated using interatomic distances and a list of atoms with Van der Waals parameters. |
| distance<br>[1], float | Interatomic distance between atoms in Å, computed from the xyz atomic coordinates taken from the PDB file. |

### EGNN

The PDB files were converted into protein-protein interfaces (PPI) in the form of residue-level graphs. For a given PDB file representing a pMHC complex, we defined the interface graph as all the residues which carbon α atoms fall within a 10 Å radius of any peptide’s residue carbon α. The only features that were used for residues are the residue type, coordinates, and a binary label representing whether the residue comes from the peptide or the MHC. Edges were defined using a k-nearest neighbors (KNN) procedure, where a node is connected to 10 of its closest neighbors. A binary edge attribute distinguishes between cross-chain edges and inter-chain edges. Node and edge features used are summarized in **Table 3**. The EGNN network is trained in a supervised manner and a self-supervised manner (i.e., 3D-SSL). The details are described in **Supplementary Note 2**.

**Table 3.** Input features to the EGNN architecture.

| Feature & Shape & Type | Description |
| --- | --- |
| NODE FEATURES |  |
| res_type<br>[1], int | Index of node residue type. |
| pos<br>[3], float | The xyz coordinates of the residue C-alpha. |
| entity<br>[1], bool | Boolean indicating whether the node is the peptide (1) or the protein (0). Used for pooling. |
| EDGE FEATURES |  |
| same_chain<br>[1], bool | Boolean indicating whether the edge connects nodes belonging to the same chain (1) or separate chains (0). |

### Sequence-based methods

The original script from the MHCFlurry 2.0 work^11^ was used to build a combined ensemble made of several neural networks. For each data configuration, the only parameters modified from the MHCflurry 2.0 script were the data input and output paths. The original script is available on https://github.com/openvax/mhcflurry/blob/master/downloads-generation/models_class1_pan/GENERATE.sh. For further details see **Supplementary Note 3**.

The MLP we used consists of a simple architecture built using PyTorch 2.13 and it is described in detail in **Supplementary Note 3**.

### HBV case study

Immunopeptidomics data was obtained by liquid chromatography tandem mass spectrometry (LC-MS/MS) as described in Kessler *et al*.^42^. In short, monocyte-derived dendritic cells (moDCs) from 8 healthy donors were pulsed with 12 prototype hepatitis B virus (HBV) synthetic long peptides (SLPs) in combination with a maturation stimulus for 22 hours. HLA-I-peptide complexes were immunoprecipitated and subjected to data-dependent acquisition LC-MS/MS analysis. Data was analyzed in PEAKS with FDR5. Obtained spectra from cross-presented, SLP-derived peptides were manually validated by two MS experts. For the values reported in **Fig. 4B**, NetMHCpan 4.1b from the web server^10^ and MHCflurry2.0 locally installed^11^ where used to predict the peptides’ binding to HLA-A*02:01 with default options.

## Data & Code Availability

Binding affinity data, 3D models, allele clusters and the trained networks will be made available on zenodo with DOIs upon acceptance. The code for data generation, featurization and training is available at https://github.com/DeepRank/3D-Vac.

## Authors Contribution

Conceived the study: LX, DFM. Prepared the data: DFM, DL, HS. 3D-CNN: DFM, DTR, HS. GNN: GC, CL. EGNN: TR. 3D-SSL: TR. MHCFlurry and MLP: DL, DFM. HBV data collection: ALK, SIB. Project supervision and coordination: LX, EB. Manuscript writing and editing: DFM, GC, LX, SG, PL, DB, DL, ALK, SIB, EB. Figures: DFM, GC, LX, NR.

All authors read and contributed to the manuscript. Giulia and Dario agree that the order of their respective names may be changed for personal pursuits to best suit their own interests.

## Acknowledgments

LX and DM are supported by the Hypatia Fellowship from Radboudumc (Rv819.52706), Hanarth Fond and the Kika grant (grant number 454). GC and SG are supported by the Netherlands eScience Center under grant number NLESC.OEC.2021.008. This work was also supported by the NVIDIA Academic Hardware Grant Program. We thank SurfSara for their generous GPU and CPU supercomputing grants (EINF-2380 and EINF-4578). We sincerely thank the funding through the AI-For-Health program at Radboudumc for H. Severin and D. Lepikov. We thank the Erasmus fellowship to D. Lepikov. We also thank Farzaneh Meimandi Parizi and Coos Baakman, the main developers of the preprocessing part of the DeepRank2 package, and Sven van der Burg, who helped with the refactoring and the maintenance of the package in the first phase of the project. We thank Prof. Peter-Bram ‘t Hoen for the constructive scientific discussions and advice.

