## Supplementary Notes for "Improving generalizability for MHC-I binding peptide predictions through geometric deep learning"

### Supplementary Note 1 - Performances in terms of AUPRC

We have evaluated our models' performances in terms of Area Under the Precision-Recall Curve (AUPRC), which is more recommended for datasets with imbalance class distributions than the AUC[1]. The AUPRC values are in line with the AUC results shown in the main text, and confirm that our StrB methods maintain better performance in the allele-clustered experiment (**Suppl. Fig. 1B**)

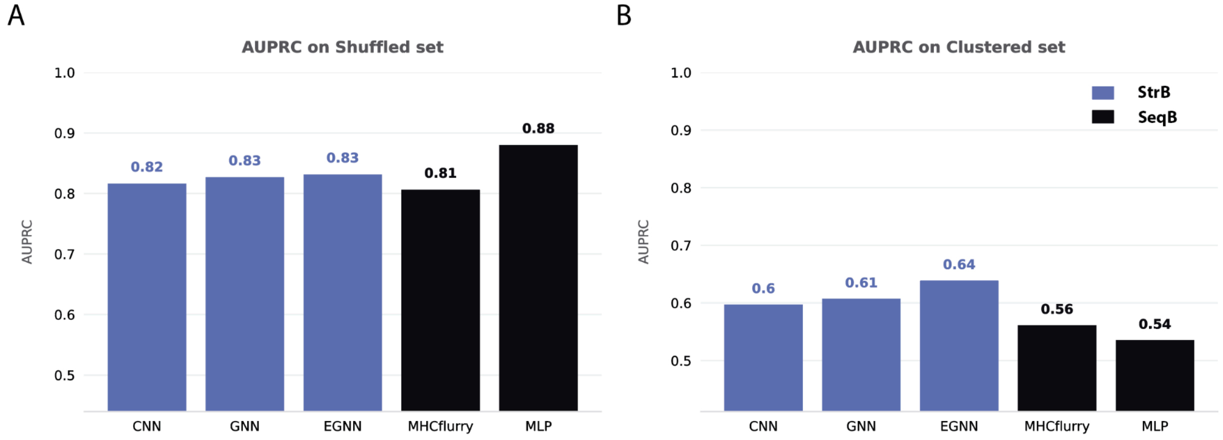


**Supplementary Figure 1. AUPRC values of StrB and SeqB methods on shuffled and allele-clustered sets.** A) StrB and SeqB methods AUPRC on the shuffled test dataset. Y-axis starting at the AUCPR baseline value for this test set of 0.44. B) StrB and SeqB methods AUPRC on the clustered-allele test dataset. Y-axis starting at the AUCPR baseline value for this test set of 0.41.

### Supplementary Note 2 - Our StrB approaches

#### 3D-CNN network architecture

The 3D-CNN architecture consists of two projection layers, two convolution layers and a three-layer MLP. First, a batchnorm layer is applied to normalize the feature according to the dataset average. Then, two 3D convolutional layers of kernel size 1 are applied as projection layers, each followed by another batchnorm layer and a ReLU activation function. Two other layers of 3D convolution with kernel size 3 are then applied, followed by a batchnorm and a max pooling layer by a factor of 2 in each dimension, in order to reduce the dimensionality of the input. The data is then flattened, piped into a batchnorm layer and fed into a small MLP of three linear layers, ending in two output neurons (one for each class between binder/non binder), followed by a Softmax for normalization. The network is trained over 15 epochs, using an Adam optimizer with a learning rate of $1e^{-3}$, a batch size of 128 and a simple cross entropy loss function.

CNN Architecture Summary:

----------------------------------------------------------------

| Layer (type) | Output Shape | Param # |
|---|---|---|
| BatchNorm3d-1 | [-1, 24, 35, 30, 30] | 48 |
| Conv3d-2 | [-1, 12, 35, 30, 30] | 300 |
| BatchNorm3d-3 | [-1, 12, 35, 30, 30] | 24 |
| ReLU-4 | [-1, 12, 35, 30, 30] | 0 |
| Conv3d-5 | [-1, 12, 35, 30, 30] | 156 |
| BatchNorm3d-6 | [-1, 12, 35, 30, 30] | 24 |
| ReLU-7 | [-1, 12, 35, 30, 30] | 0 |
| Conv3d-8 | [-1, 24, 33, 28, 28] | 7,800 |
| BatchNorm3d-9 | [-1, 24, 33, 28, 28] | 48 |
| MaxPool3d-10 | [-1, 24, 16, 14, 14] | 0 |
| ReLU-11 | [-1, 24, 16, 14, 14] | 0 |
| Conv3d-12 | [-1, 48, 14, 12, 12] | 31,152 |
| BatchNorm3d-13 | [-1, 48, 14, 12, 12] | 96 |
| MaxPool3d-14 | [-1, 48, 7, 6, 6] | 0 |
| ReLU-15 | [-1, 48, 7, 6, 6] | 0 |
| Flatten-16 | [-1, 12096] | 0 |
| BatchNorm1d-17 | [-1, 12096] | 24,192 |
| Linear-18 | [-1, 128] | 1,548,416 |
| ReLU-19 | [-1, 128] | 0 |
| Dropout-20 | [-1, 128] | 0 |
| Linear-21 | [-1, 128] | 16,512 |
| ReLU-22 | [-1, 128] | 0 |
| Dropout-23 | [-1, 128] | 0 |
| Linear-24 | [-1, 2] | 258 |
| Softmax-25 | [-1, 2] | 0 |

================================================================

Total params: 1,629,026
Trainable params: 1,629,026
Non-trainable params: 0

----------------------------------------------------------------

Input size (MB): 2.88
Forward/backward pass size (MB): 35.55
Params size (MB): 6.21
Estimated Total Size (MB): 44.64

----------------------------------------------------------------

**GNN network architecture**

The structure of the GNN architecture consists first of three message passing layers on the top of each other, which have been designed for aggregating information from neighboring nodes and edges. The message passing framework[2] is an ubiquitous paradigm for designing GNNs for molecular data. It involves the iterative update of node representations by aggregating information from neighboring nodes. Here, we defined each message passing layer as a linear layer followed by the ReLu activation function, that takes as input a concatenation of edges' and neighborhood nodes' features, and gives as output messages of size 32 (message function) which are then aggregated using a sum function as aggregation function. Then the node features concatenated with the aggregated messages are passed through the so-called update function, represented here by a linear layer followed by the ReLu activation function. The output from the third message passing layer is then passed through a readout aggregation function that performs the mean, and finally three sequential linear layers of output size 128, the first two followed by the ReLU activation function, are applied. Finally, the output is normalized by the SoftMax activation function and represents the probability between 0 and 1 for each binary BA class. During the training we used Adam optimizer with a learning rate of 0.001 and a batch size of 64. An early stopping regularization monitored the performance of the network on the validation set and stopped the training process at epoch 34 out of 70.

GNN Architecture summary:

```
=================================================================
Layer (type:depth-idx)                   Param #
=================================================================
├─MessagePassingLayer: 1-1               --
│    └─Sequential: 2-1                   --
│    │    └─Linear: 3-1                  3,840
│    │    └─ReLU: 3-2                    --
│    └─Sequential: 2-2                   --
│    │    └─Linear: 3-3                  5,130
│    │    └─ReLU: 3-4                    --
├─MessagePassingLayer: 1-2               --
│    └─Sequential: 2-3                   --
│    │    └─Linear: 3-5                  3,840
│    │    └─ReLU: 3-6                    --
│    └─Sequential: 2-4                   --
│    │    └─Linear: 3-7                  5,130
│    │    └─ReLU: 3-8                    --
├─MessagePassingLayer: 1-3               --
│    └─Sequential: 2-5                   --
│    │    └─Linear: 3-9                  3,840
│    │    └─ReLU: 3-10                   --
│    └─Sequential: 2-6                   --
│    │    └─Linear: 3-11                 5,130
│    │    └─ReLU: 3-12                   --
├─Sequential: 1-4                        --
│    └─Linear: 2-7                       7,424
│    └─ReLU: 2-8                         --
│    └─Linear: 2-9                       16,512
│    └─ReLU: 2-10                        --
│    └─Linear: 2-11                      258
=================================================================
Total params: 51,104
```

Trainable params: 51,104
Non-trainable params: 0
==============================================================

### EGNN network architecture

The EGNN architecture used is a modified version of the E(n)-Equivariant Graph Neural Network[3]. Following the standard message-passing formula, its message, aggregate and update function are realized with the following four equations:

$$m_{ij} = MLP_{msg}(h_i^l, h_j^l, RBF(\|x_i^l - x_j^l\|), c_{ij})$$

$$x_i^{l+1} = x_i^l + \frac{1}{|N(i)|}\sum_{j\neq i}(x_i^l - x_j^l)MLP_{pos}(m_{ij})$$

$$m_i = \sum_{j\neq i} m_{ij}$$

$$h_i^{l+1} = MLP_{upd}(h_i^l, m_i)$$

Where $h_i^l$ is the embedding of node $i$ at layer $l$. $x_i^l$ is the coordinate of CA atom of the residue $i$ at layer $l$. $c_{ij}$ is the edge features between nodes $i$ and $j$. The message function of our network is a feed-forward NN (MLP) which takes in the representations of two nodes, alongside a radial basis function (RBF) embedding of the distance between them and the binary edge type.

Notably different from vanilla GNN architectures, our network includes an equivariant position update step, ensuring the propagation of a vector quantity through its layers, which in turns allows for greater distinguishing power of graph geometry[3]. Our implementation differs from the implementation of EGNN in Satorras *et al.*[3] in two regards. Firstly, we use a RBF embedding of the distance between nodes $r_{ij}$, as is common in modern equivariant networks[4,5]. Gaussian RBFs are functions of distance which take the form:

$$\phi_k(r_{ij}) = exp(-\frac{(r_{ij}-\mu_k)^2}{2\sigma_k^2})$$

We use a 64-dimensional RBF encoding of distances, with the $u_k$ evenly distributed between 0 and 30Å, and a $\sigma_k$ value of 1.

Secondly, we implement the message MLP using strong-conditioned linear layers[6], ensuring a more expressive conditioning of the message on the distance between nodes. A common way of conditioning on distances in linear layers is to concatenate the distance to other input features, called by Koishekenov and Bekkers[6] "weak conditioning". We opt instead for strong conditioning, which has been shown to outperform its weak counterpart. In strong conditioning, a linear layer is parametrized by two matrices $W^h$ and $W^d$ . It operates on a feature vector `*h*` and a conditioning vector `*d*` as follows:

$$cond\_linear(h, d) = W^h h \, ⍰ \, W^d d$$

where ⍰ is elementwise multiplication.

The EGNN network is composed of three such graph layers, following an initial embedding of the node residue type, which is represented as integers. The node embedding dimension is 128. MLP_msg, MLP_pos, MLP_upd have two layers, and use the ReLU activation layer and layer normalization. The binary edge feature ('Is_same_chain') is concatenated to the node features before being passed through the message MLP. We optimized all EGNNs using the Adam optimizer, with the learning rate and weight decay set to $1e - 4$ .

This EGNN is used for both the supervised case and the self-supervised case.
A. *Supervised case (EGNN).* In the supervised case, the binding label is formed as a sum of individual peptide residue contributions, formed by passing the residue embeddings (size/node: 128) through a two-layer residue-wise MLP (architecture: 128 ->128 -> 1, activation function: ReLU). This sum is then passed through a sigmoid layer to ensure an output between 0 and 1. Loss function is cross entropy. For the supervised experiment, we use a batch size of 512.

*B. Self-supervised case (3D-SSL).* In the SSL case, the residue embedding (size/node: 128) is passed to a two-layer MLP (architecture: 128 -> 128 -> 23, activation function: ReLU). The Softmax is applied to the output of this two-layer MLP to return a distribution over residue types (i.e., a vector with size of 23 by 1, 22 residue types + UNK/MASK). We use cross entropy as the loss function. During training, a random 20% of residues are masked. For SSL training, we use a batch size of 128.

Once the SSL network is trained, we use it to provide scores for PANDORA models. The score is a sum of individual peptide residue contributions to the binding. Residue contributions are calculated in the following manner:

1. First, the i-th residue is masked, as in the training procedure. The network predicts the probability of the 20 types of amino acids of the masked residue given its structural neighborhood, $P(A_i)$ where $A_i$ is the amino acid type, $A_i \in \{ALA, ARG, ...\}$.
2. Convert this probability $P(A_i)$ into statistical potentials using Boltzmann Distribution:

$$U(\vec{x}_i) = - KT * logP(A_i) - KT * logP(A_i)_{ref}$$

where $U(\vec{x}_i)$ is the energy contribution of residue $i$, $\vec{x}_i$ is the (x,y,z) coordinates of the masked amino acid, $K$ is Boltzmann constant, and $T$ is the temperature. In our current implementation, we ignored the reference state (left for future studies). And we chose $KT = 1$ without losing the generalizability.

The above two steps are repeated for every residue in the peptide, with the resulting score being a sum of the individual log-probabilities. Note that the procedure requires n forward passes of the MRP network for a peptide n-mer. If the statistical potential learned by the network is accurate for pMHC complexes, it should correlate well with the peptide binding affinity.

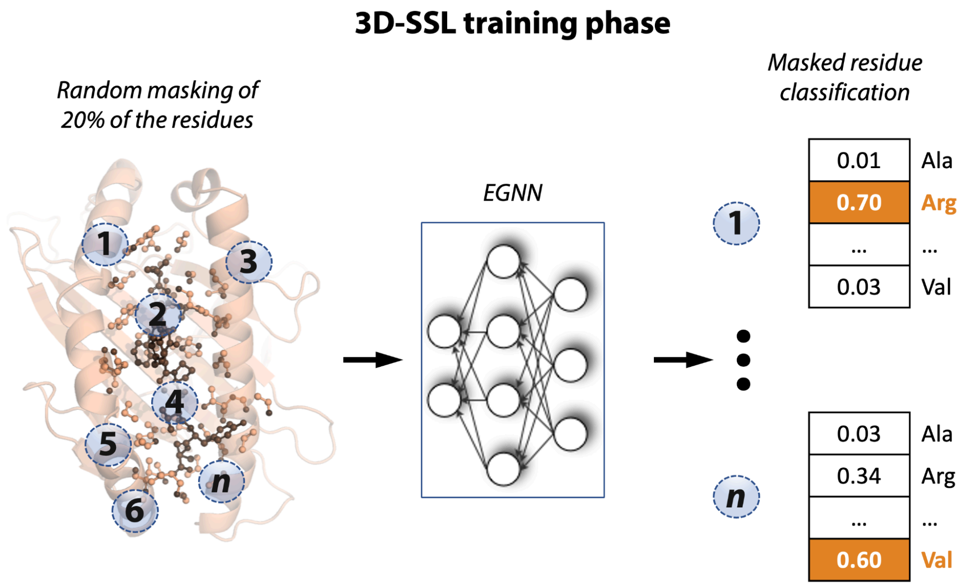


**Supplementary Figure 2. 3D-SSL training phase.** During the training phase of 3D-SSL, each 3D structure is converted into a residue-level graph, and 20% of its nodes are randomly masked. The graph is then fed into EGNN, which returns a residue identity probability for each masked node.

EGNN Architecture summary:

| Layer | Input Shape | Output Shape | #Param |
|---|---|---|---|
| EGNNModel | [61, 61] | [2, 1] | 443,969 |
| ⊢(embedding)AtomEmbedding | [61, 61] | [61, 128] | 2,944 |
| └(embedding)Embedding | [61] | [61, 128] | 2,944 |
| ⊢(convs)ModuleList | -- | -- | 424,128 |
| └(0)EGNNLayer | [61, 128],[61, 3],[2, 718],[718, 1] | [61, 128],[61, 3] | 141,376 |
| └(1)EGNNLayer | [61, 128],[61, 3],[2, 718],[718, 1] | [61, 128],[61, 3] | 141,376 |
| └(2)EGNNLayer | [61, 128],[61, 3],[2, 718],[718, 1] | [61, 128],[61, 3] | 141,376 |
| ⊢(pred)GraphLevelPredictionHead | [61,128], [61,61] | [2, 1] | 16,897 |
| └(pred)LinearHead | [9,128] | [9, 1] | 16,897 |
| └(activation)ReLU | [9,128] | [9, 128] | -- |
| └(mlp)Sequential | [9,128] | [9, 1] | 16,897 |

### Supplementary Note 3 - SeqB approaches

**Input for MLP and MHCflurry** (MHC pseudoseq + peptide 45-mer representation)

Both the MLP and the re-trained MHCFlurry 2.0[7] sequence-based predictors used the same BLOSUM62 substitution matrix[8] as amino acids representation. Inspired by the MHCflurry 2.0 encoding method, the MHC allele and peptide are encoded differently and concatenated together: a) the MHC allele is encoded by a fixed length of 37 binding groove residues known to interact with the peptide and called "pseudosequence", as originally introduced by Hoof *et al.*[9] b) The peptide is encoded as in O'Donnell *et al.*[7], as a concatenated array of three representations of maximum length of 15: left-aligned, centered and right-aligned. For peptides having less than 15 residues, the three representations are supplemented with the placeholder residue X to reach the maximum length of the peptide. Assuming the 5-residues peptide

AVFGK, AVFGKXXXXXXXXXX, XXXXXAVFGKXXXXX XXXXXXXXXXAVFGK are its left-aligned, centered and right-aligned representations, respectively. The final input for the sequence-based methods is a concatenated peptide-MHC matrix of size 82*21 (45*21 for the peptide + 37*21 for the MHC pseudosequence).

**MLP architecture**

The MLP is made of a BatchNorm1d() input layer followed by 3 hidden layers of 512 neurons. Each hidden layer has a dropout rate of 50%. The optimizer used was Adam with binary cross entropy as loss function.

**MHCFlurry retraining**

Scripts from the MHCflurry 2.0[10] github release were used to build a combined ensemble made of several models. For each experiment (shuffled and allele-clustered dataset), the only parameters modified from the MHCflurry 2.0 script used to generate the model were the data input and output paths.
The original script is available at https://github.com/openvax/mhcflurry/blob/master/downloads-generation/models_class1_pan/GENERATE.sh.

We note that the performance of our retrained version of MHCflurry 2.0 is lower in our study than previously reported[10]. One possible factor causing this reduced performance is the considerably smaller amount of training data we used in this proof-of-concept study compared to the original publication. In addition, its performance could be negatively impacted by having used the BA predictor only, while the final score of MHCflurry 2.0 reported in O'Donnell *et al.*[10] is a combination of the single scores from the BA and the antigen processing (AP) predictors. We did not include the AP predictor because we focused on predictions regarding MHC allele-dependent effects only (BA prediction), and not allele-independent effects (AP prediction). Second, MHCflurry 2.0 explorable hyperparameters during learning were designed on a much larger dataset, and not optimized for the restricted one we used. Lastly, we made a deliberate choice to omit the pre-training step of MHCflurry 2.0, in which the network is initially trained on several hundred million synthetic data labeled with MHCFlurry 1.2.0 and subsequently fine-tuned on the actual data. We opted against this step due to concerns that utilizing randomly generated peptides pre-labeled by another model might cause a data leakage from the initial model to the second one, compromising the intended separation of our data.

### Supplementary Note 4 - Computational efficiency of three StrB methods

Regarding the computational costs of our three StrB approaches, the GNN architecture demonstrated results comparable to CNN in terms of performance metrics, but has clear computational efficiency advantages: performance maintained with a coarser-grained resolution (residue-level graphs vs. atomic-level grids), faster featurization and data mapping, and lower storage requirements. Notably, the EGNN showed the best predictive power, and with even faster data mapping and featurization and lower storage requirement (**Suppl. Table 1**).

**Supplementary Table 1. Computational costs of our StrB supervised methods.** Our EGNN showed the best prediction performance, with fastest data mapping and featurization and lowest storage requirement.

| | featurization and data mapping on one CPU core/pMHC model | hard-drive storage/pMHC model |
|---|---|---|
| 3D-CNN | 3.2±0.1sec | 744±17 KB |
| GNN | 6.2±0.1sec | 5.6±0.2 MB |
| EGNN | 0.07±0.02 sec | 3.9±0.1 KB |
